# Dopamine, Inference, and Uncertainty

**DOI:** 10.1101/149849

**Authors:** Samuel J. Gershman

## Abstract

The hypothesis that the phasic dopamine response reports a reward prediction error has become deeply entrenched. However, dopamine neurons exhibit several notable deviations from this hypothesis. A coherent explanation for these deviations can be obtained by analyzing the dopamine response in terms of Bayesian reinforcement learning. The key idea is that prediction errors are modulated by probabilistic beliefs about the relationship between cues and outcomes, updated through Bayesian inference. This account can explain dopamine responses to inferred value in sensory preconditioning, the effects of cue pre-exposure (latent inhibition) and adaptive coding of prediction errors when rewards vary across orders of magnitude. We further postulate that orbitofrontal cortex transforms the stimulus representation through recurrent dynamics, such that a simple error-driven learning rule operating on the transformed representation can implement the Bayesian reinforcement learning update.

## 1 Introduction

The phasic firing of dopamine neurons in the midbrain has long been thought to report a reward prediction error—the discrepancy between observed and expected reward—whose purpose is to correct future reward predictions (Eshel et al., 2015; Glimcher, 2011; Schultz et al., 1997). This hypothesis can explain many key properties of dopamine, such as its sensitivity to the probability, magnitude and timing of reward, its dynamics over the course of a trial, and its causal role in learning. Despite its success, the prediction error hypothesis faces a number of puzzles. First, why do dopamine neurons respond under some conditions, such as sensory preconditioning (Sadacca et al., 2016) and latent inhibition (Young et al., 1993), where the prediction error should theoretically be zero? Second, why do dopamine responses appear to rescale with the range or variance of rewards (Tobler et al., 2005)? These phenomena appear to require a dramatic departure from the normative foundations of reinforcement learning that originally motivated the prediction error hypothesis (Sutton and Barto, 1998).

This paper provides a unified account of these phenomena, expanding the prediction error hypothesis in a new direction while retaining its normative foundations. The first step is to reconsider the computational problem being solved by the dopamine system: instead of computing a single point estimate of expected future reward, the dopamine system recognizes its own uncertainty by computing a probability distribution over expected future reward. This probability distribution is updated dynamically using Bayesian inference, and the resulting learning equations retain the important features of earlier dopamine models. Crucially, the Bayesian theory goes beyond earlier models by explaining why dopamine responses are sensitive to sensory preconditioning, latent inhibition and reward variance.

The theory presented here was first developed to explain a broad range of associative learning phenomena within a unifying framework (Gershman, 2015). We extend this theory further by equipping it with a mechanism for updating beliefs about cue-specific volatility (i.e., how quickly associations between particular cues and outcomes change over time). This mechanism harkens back to the classic Pearce-Hall theory of attention in associative learning (Pearce and Hall, 1980; Pearce et al., 1982), as well as to more recent Bayesian incarnations (Behrens et al., 2007; Mathys et al., 2011; Nassar et al., 2010; Yu and Dayan, 2005). As shown below, volatility estimation is important for understanding the effect of reward variance on the dopamine response.

## 2 Temporal difference learning

The prediction error hypothesis of dopamine was originally formalized by Schultz et al. (1997) in terms of the temporal difference (TD) learning algorithm (Sutton and Barto, 1998), which posits an error signal of the form:

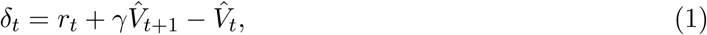

where *r_t_* is the reward received at time *t, γ ∊* [0, 1] is a discount factor that down-weights distal rewards exponentially, and 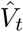 is an estimate of the expected discounted future return (or *value):*

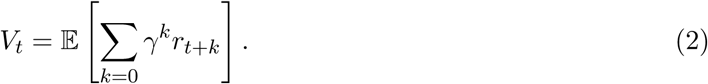

By incrementally adjusting the parameters of the value function to minimize prediction errors, TD learning gradually improves its estimates of future rewards. One common functional form, both in machine learning and neurobiological applications, is linear function approximation, which approximates the value as a linear function of features 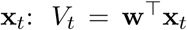, where **w** is a vector of weights, updated according to 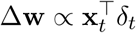.

In the complete serial compound (CSC) representation, each cue is broken down into a cascade of temporal elements, such that each feature corresponds to a binary variable indicating whether a stimulus is present or absent at a particular point in time. This allows the model to generate temporally precise predictions, which have been systematically compared to phasic dopamine signals. While the original work by Schultz et al. (1997) showed good agreement between prediction errors and dopamine using the CSC, later work called into question its adequacy (Daw et al., 2006; Gershman et al., 2014; Ludvig et al., 2008). Nonetheless, we will adopt this representation for its simplicity, noting that our substantive conclusions are unlikely to be changed with other temporal representations.

## 3 Reinforcement learning as Bayesian inference

The TD model is a point estimation algorithm, updating a single weight vector over time. Gershman (2015) argued that associative learning is better modeled as Bayesian inference, where a probability distribution over all possible weight vectors is updated over time. This idea was originally explored by Dayan and colleagues (Dayan and Kakade, 2001; Dayan et al., 2000) using a simple Bayesian extension of the Rescorla-Wagner model (the Kalman filter). This model can explain retrospective revaluation phenomena like backward blocking that posed notorious difficulties for classical models of associative learning (Miller et al., 1995). Gershman (2015) illustrated the explanatory range of the Kalman filter by applying it to numerous other phenomena. However, the Kalman filter is still fundamentally limited by the fact that it is a trial-level model and hence cannot explain the effects of intra-trial structure like the interstimulus interval or stimulus duration. It was precisely this structure that motivated real-time frameworks like the TD model (Sutton and Barto, 1990).

The same logic that transforms Rescorla-Wagner into the Kalman filter can be applied to transform the TD model into a Bayesian model (Geist and Pietquin, 2010). Gershman (2015) showed how the resulting unified model (Kalman TD) can explain a range of phenomena that neither the Kalman filter nor the TD model can explain in isolation. In this section, we describe Kalman TD and its extension to incorporate volatility estimation. We then turn to studies of the dopamine system, showing how the same model can provide a more complete account of dopaminergic prediction errors.

### 3.1 Kalman temporal difference learning

To derive a Bayesian model, we first need to specify the data-generating process. In the context of associative learning, this entails a prior probability distribution on the weight vector, *p*(**w**_0_), a *change process* on the weights, *p*(**w***_t_|***w***_t_*_−1_), and a reward distribution given stimuli and weights, *p*(*r_t_|***w***_t_*, **x***_t_*). Kalman TD assumes a linear-Gaussian dynamical system:

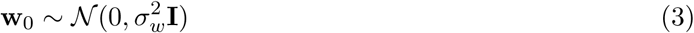

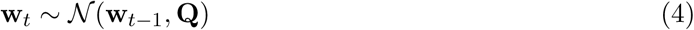

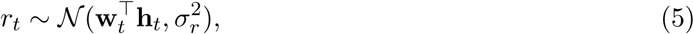

where **I** is the identity matrix, **Q** = diag(*q*_1_,…,*q_D_*) is a diagonal diffusion covariance matrix, and **h***_t_* = *γx_t+_*_1_– **x***_t_* is the discounted temporal derivative of the features. Under these assumptions, the value function can be expressed as a linear function of the features, *V_t_* = **w^T^x***_t_*, just as in the linear function approximation architecture for TD (though in this case the relationship is exact).

Given an observed sequence of feature vectors, Bayes’ rule stipulates how to compute the posterior distribution over the weight vector:

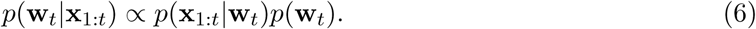

Under the generative assumptions described above, the Kalman filter can be used to update the parameters of the posterior distribution in closed form. In particular, the posterior is Gaussian with mean **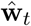** and covariance matrix 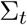, updated according to:

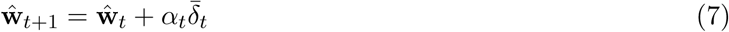

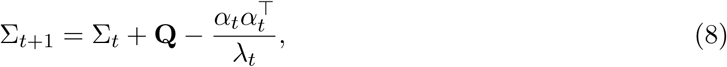

where **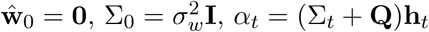** is a vector of learning rates, and

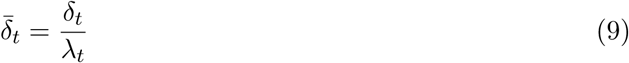

is the prediction error rescaled by the marginal variance

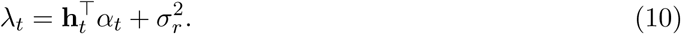

Like the original TD model, the Kalman TD model posits updating of weights by prediction error, and the core empirical foundation of the TD model (see Glimcher, 2011) also applies to Kalman TD. Unlike the original TD model, the learning rates change dynamically in Kalman TD, a property important for explaining phenomena like latent inhibition, as discussed below. Another deviation from the original TD model is the fact that weight updates may not be independent across cues; if there is non-zero covariance between cues, then observing novel information about one cue will change beliefs about the other cue. This property is instrumental to the explanation of various revaluation phenomena (Gershman, 2015), which we explore below using the sensory preconditioning paradigm.

### 3.2 Volatility estimation

One unsatisfying, counterintuitive property of the Kalman TD model is that the learning rates do not depend on the reward history. This means that the model will not be able to capture changes in learning rate due to variability in the reward history. There is in fact a considerable literature suggesting learning rate changes as a function of reward history, though the precise nature of such changes is controversial (Le Pelley, 2004; Mitchell and Le Pelley, 2010). For example, learning is slower when the cue was previously a reliable predictor of reward (Hall and Pearce, 1979). Pearce and Hall (1980) interpreted this and other findings as evidence that learning rate declines with cue-outcome reliability. They formalized this idea by assuming that learning rate is proportional to the absolute prediction error (see Roesch et al., 2012, for a review of the behavioral and neural evidence).

A similar mechanism can be derived from first principles within the Kalman TD framework. We have so far assumed that the diffusion covariance matrix **Q** is known, but the diagonal elements (diffusion variances) *q*_1_,…, *q_D_* can be estimated by maximum likelihood methods. In particular, we can derive a stochastic gradient descent rule by differentiating the log-likelihood with respect to the diffusion variances:

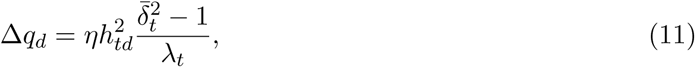

where *η* is a learning rate. This update rule exhibits the desired dependence of learning rate on the unsigned prediction error (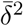), since learning rate for cue *d* increases monotonically with *qd.*

### 3.3 Modeling details

We use the same parameters as in our earlier paper (Gershman, 2015): 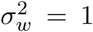, 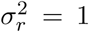, and *γ* = 0.98. The volatilities were initialized to *qd* = 0.01 and then updated using a meta-learning rate of *η* = 0.1. Stimuli were modeled with a 4-time-step CSC representation and an intertrial interval of 6 time-steps.

## 4 Applications to the dopamine system

We are now in a position to resolve the puzzles with which we started, focusing on two empirical implications of the TD model. First, the model only updates the weights of present cues, and hence cues that have not been paired directly with reward or with reward-predicting cues should not elicit a dopamine response. This implication disagrees with findings from a sensory preconditioning procedure (Sadacca et al., 2016), where cue A is sequentially paired with cue B and cue C is sequentially paired with cue D (Figure 1). If cue B is subsequently paired with reward and cue D is paired with nothing, cue A comes to elicit both a conditioned response and elevated dopamine activity compared to cue B. The TD model predicts no dopamine response to either A or B. The Kalman TD model, in contrast, learns a positive covariance between the sequentially presentedcues. As a consequence, the learning rates will be positive for both cues whenever one of them is presented alone, and hence conditioning one cue in a pair will cause the other cue to inherit value.

**Figure 1:**
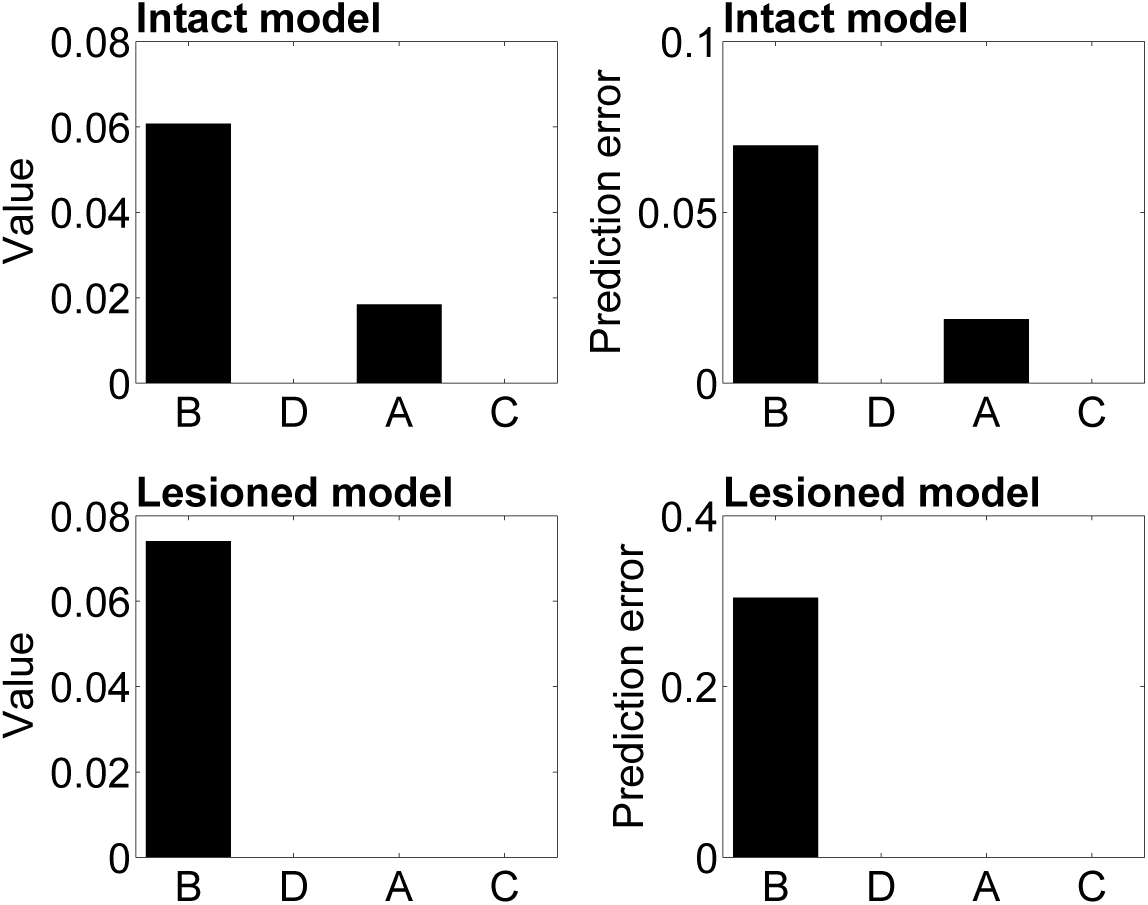
Sensory preconditioning: simulation of Sadacca et al. (2016). In the sensory preconditioning phase, animals are exposed to the cue sequences A→B and C→D. In the conditioning phase, B is associated with reward, and D is associated with nothing. The intact OFC model shows elevated conditioned responding and dopamine activity when subsequently tested on cue A, but not when tested on cue C. The lesioned OFC model does not show this effect, but responding to B is intact.

**Figure 2:**
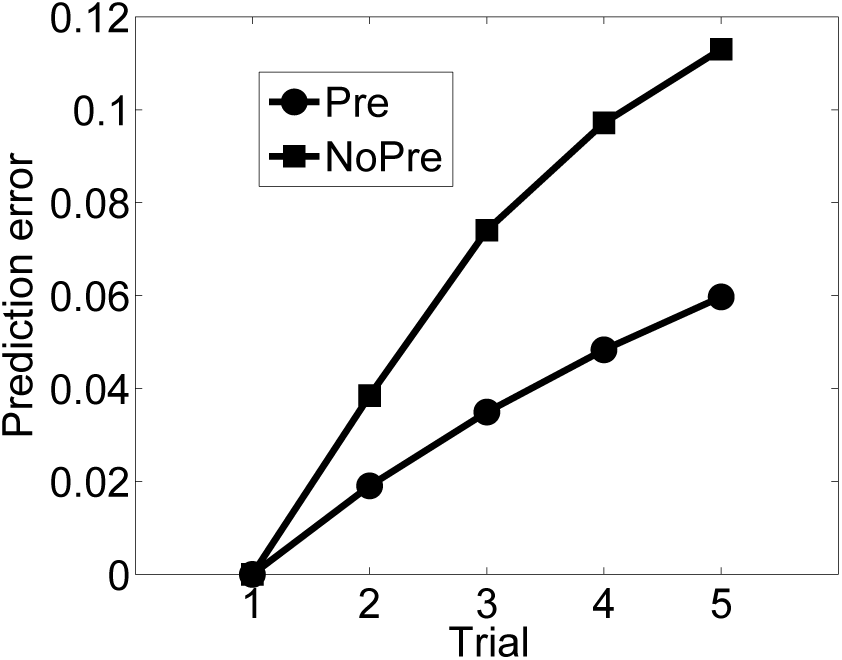
Latent inhibition: simulation of Young et al. (1993). The acquisition of a dopamine response to a cue is slower for pre-exposed cues (Pre) compared to novel cues (NoPre).

In the latent inhibition procedure, pre-exposing a cue prior to pairing it with reward should have no effect on dopamine responses during conditioning in the original TD model (since the prediction error is 0 throughout pre-exposure), but experiments show that pre-exposure results in a pronounced decrease in both conditioned responding and dopamine activity during conditioning (Young et al., 1993). The Kalman TD model predicts that the posterior variance will decrease with repeated pre-exposure presentations (Gershman, 2015) and hence the learning rate will decrease as well. This means that the prediction error signal will propagate more slowly to the cue onset for the pre-exposed cue compared to the non-pre-exposed cue.

A second implication of the TD model is that dopamine responses at the time of reward should scale with reward magnitude. This implication disagrees with the work of (Tobler et al., 2005), who paired different cues half the time with a cue-specific reward magnitude (liquid volume) and half the time with no reward. Although dopamine neurons increased their firing rate whenever reward was delivered, the size of this increase was essentially unchanged across cues despite the reward magnitudes varying over an order of magnitude. Tobler et al. (2005) interpreted this finding as evidence for a form of “adaptive coding,” whereby dopamine neurons adjust their dynamic range to accommodate different distributions of prediction errors (see also Diederen and Schultz, 2015; Diederen et al., 2016, for converging evidence from humans). Adaptive coding has been found throughout sensory areas as well as in reward-processing areas (Louie and Glimcher, 2012). While adaptive coding can be motivated by information-theoretic arguments (Atick, 1992), the question is how to reconcile this property with the TD model.

The Kalman TD model resolves this puzzle if one views dopamine as reporting 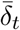 (the variance-scaled prediction error) instead of *δ_t_* (Figure 3).^1^ Critical to this explanation is volatility updating: the scaling term (*λ_t_*) increases with the diffusion variance *q_d_*, which itself scales with the reward magnitude in the experiment of Tobler et al. (2005). In the absence of volatility updating, diffusion variance would stay fixed, and hence *λ_t_* would no longer be a function of reward history.^2^

**Figure 3:**
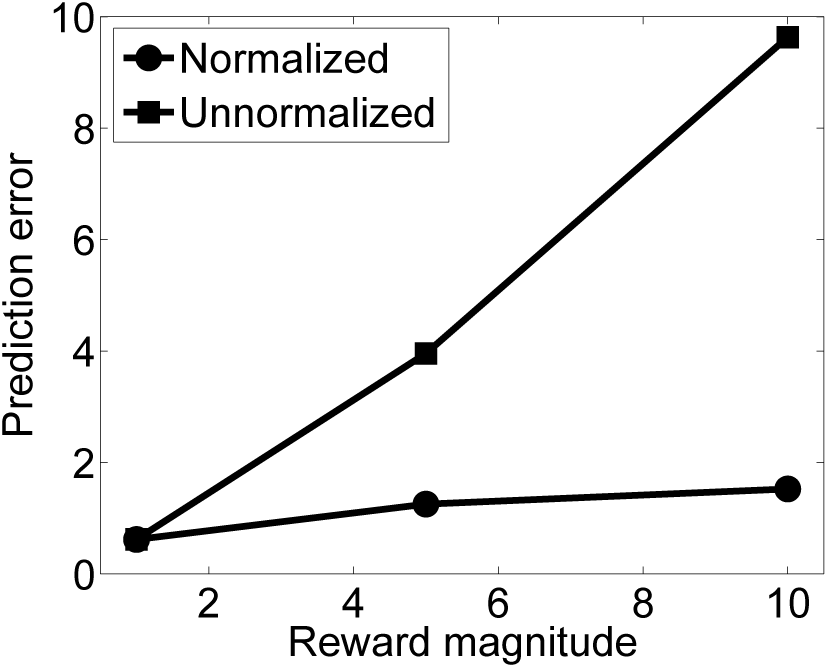
Adaptive coding: simulation of Tobler et al. (2005). Each cue is associated with a 50% chance of earning a fixed reward, and 50% chance of nothing. Dopamine neurons show increased responding to the reward compared to nothing, but this increase does not change across cues delivering different amounts of reward. This finding is inconsistent with the standard TD prediction error account, but is consistent with the hypothesis that prediction errors are divided by the posterior predictive variance.

## 5 Representational transformation in the orbitofrontal cortex

Dayan and Kakade (2001) described a neural circuit that approximates the Kalman filter, but did not explore its empirical implications. This section reconsiders the circuit implementation applied to the Kalman TD model, and then discusses experimental data relevant to its neural substrate.

The network architecture is shown in Figure 4. The input units represent the discounted feature derivatives, **h***_t_*, which are then passed through an identity mapping to the intermediate units **y***_t_*. The intermediate units are recurrently connected with a synaptic weight matrix **B** and undergo linear dynamics given by:

**Figure 4:**
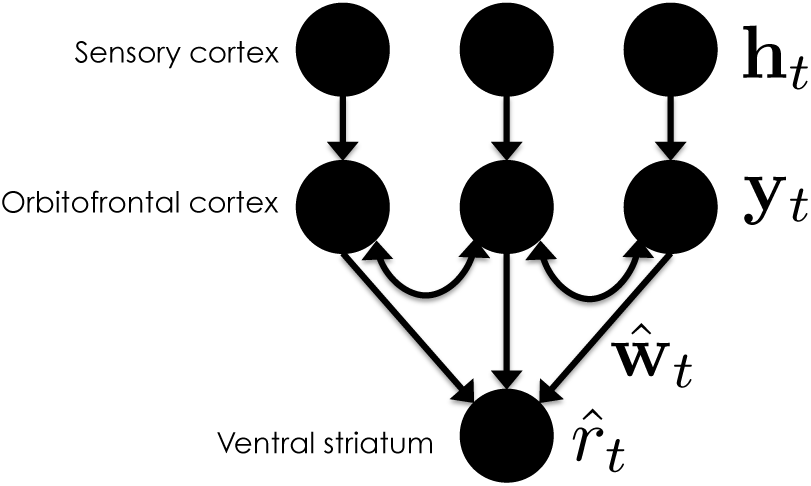
Neural architecture for Kalman TD. Modified from Dayan and Kakade (2001). Nodes corresponds to neurons, arrows correspond to synaptic connections.

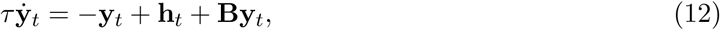

where *τ* is a time constant. These dynamics will converge to **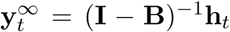**, assuming the inverse exists. The synaptic weight matrix is updated according to an anti-Hebbian learning rule (Atick and Redlich, 1993; Goodall, 1960):

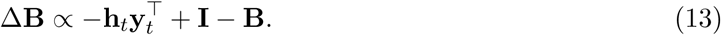

If **B** is initialized to all zeros, this learning rule asymptotically satisfies 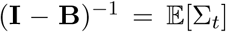. It follows that 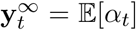 asymptotically. Thus, the intermediate units, in the limit of infinite past experience and infinite computation time, approximate the learning rates required by the Kalman filter. As noted by Dayan and Kakade (2001), the resulting outputs can be viewed as decorrelated (whitened) versions of the inputs. Instead of modulating learning rates over time (e.g., using neuromodulation; Doya, 2002; Nassar et al., 2012; Yu and Dayan, 2005), the circuit transforms the inputs so that they implicitly encode the learning rate dynamics.

The final step is to update the synaptic connections (**w**) between the intermediate units and a reward prediction unit (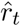):

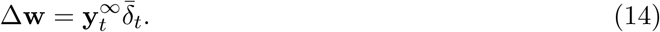

The prediction error is computed with respect to the initial output of the intermediate units (i.e., **x***_t_*), whereas the learning rates are computed with respect to their outputs after convergence. This is consistent with the assumption that the phasic dopamine response is fast, but the recurrent dynamics and synaptic updates are relatively slow.

Figure 4 presents a neuroanatomical gloss on the original proposal by Dayan and Kakade (2001). We suggest that the intermediate units correspond to the orbitofrontal cortex (OFC), with feedforward synapses to reward prediction neurons in the ventral striatum (Eblen and Graybiel, 1995). This interpretation offers a new, albeit not comprehensive, view of the OFCs role in reinforcement learning. Wilson et al. (2014) have argued that the OFC represents a “cognitive map” of task space, providing the state representation over which TD learning operates. The circuit described above can be viewed as implementing one form of state representation based on a whitening transform.

If this interpretation is correct, then OFC damage should be devastating for some kinds of associative learning (namely, those that entail non-zero covariance between cues) while leaving other kinds of learning intact (namely, those that entail uncorrelated cues). A particularly useful example of this dissociation comes from work by Jones et al. (2012), which demonstrated that OFC lesions eliminate sensory preconditioning while leaving first-order conditioning intact. This pattern is reproduced by the Kalman TD model if the intermediate units are “lesioned” such that no input transformation occurs (i.e., inputs are mapped directly to rewards; Figure 1). In other words, the lesioned model is reduced to the original TD model with fixed learning rates.

## 6 Discussion

The twin roles of Bayesian inference and reinforcement learning have a long history in animal learning theory, but until recently these ideas were not unified into a single theory known as Kalman TD (Gershman, 2015). In this paper, we applied the theory to several puzzling phenomena in the dopamine system: the sensitivity of dopamine neurons to posterior variance (latent inhibition), covariance (sensory preconditioning) and posterior predictive variance (adaptive coding). These phenomena could be explained by making two principled modifications to the prediction error hypothesis of dopamine. First, the learning rate, which drives updating of values, is vector-valued in Kalman TD, with the result that associative weights for cues can be updated even when that cue is not present, provided it has non-zero covariance with another cue. Furthermore, the learning rates can change over time, modulated by the agent’s uncertainty. Second, Kalman TD posits that dopamine neurons report a normalized prediction error, 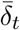, such that greater uncertainty suppresses dopamine activity (see also Preuschoff and Bossaerts, 2007).

How are the probabilistic computations of Kalman TD implemented in the brain? We modified the proposal of Dayan and Kakade (2001), according to which recurrent dynamics produce a transformation of the stimulus inputs that effectively whitens (decorrelates) them. Standard error-driven learning rules operating on the decorrelated input are then mathematically equivalent to the Kalman TD updates. One potential neural substrate for this stimulus transformation is the OFC, a critical hub for state representation in RL (Wilson et al., 2014). We showed that lesioning the OFC forces the network to fall back on a standard TD update (i.e., ignoring the covariance structure). This prevents the network from exhibiting sensory preconditioning, as has been observed experimentally (Jones et al., 2012). The idea that recurrent dynamics in OFC play an important role in stimulus representation for reinforcement learning and reward expectation has also figured in earlier models (Deco and Rolls, 2005; Frank and Claus, 2006).

Kalman TD is closely related to the hypothesis that dopaminergic prediction errors operate over belief state representations. These representations arise when an agent has uncertainty about the hidden state of the world. Bayes’ rule prescribes that this uncertainty be represented as a posterior distribution over states (the belief state), which can then feed into standard TD learning mechanisms. Several authors have proposed that belief states could explain some anomalous patterns of dopamine responses (Daw et al., 2006; Rao, 2010), and experimental evidence has recently accumulated for this proposal (Lak et al., 2017; Starkweather et al., 2017; Takahashi et al., 2016). One way to understand Kalman TD is to think of the weight vector as part of the hidden state. A similar conceptual move has been studied in computer science, in which the parameters of a Markov decision process are treated as unknown, thereby transforming it into a partially observable Markov decision process (Duff, 2002; Poupart et al., 2006). Kalman TD is a model-free counterpart to this idea, treating the parameters of the function approximator as unknown. This view allows one to contemplate more complex versions of the model proposed here, for example with nonlinear function approximators or structure learning (Gershman et al., 2015), although inference quickly becomes intractable in these cases.

A number of other authors have suggested that dopamine responses are related to Bayesian inference in various ways. Friston and colleagues have developed a theory grounded in a variational approximation of Bayesian inference, whereby dopamine reports an estimate of inverse variance (FitzGerald et al., 2015; Friston et al., 2012; Schwartenbeck et al., 2014). This theory fits well with the modulatory effects of dopamine on downstream circuits, but it is currently unclear to what extent this theoretical framework can account for the body of empirical data on which the prediction error hypothesis of dopamine is based. Other authors have suggested that dopamine is involved in specifying a prior probability distribution (Costa et al., 2015) or influencing uncertainty representation in the striatum (Mikhael and Bogacz, 2016). These different possibilities are not necessarily mutually exclusive, but more research is necessary to bridge these varied roles of dopamine in probabilistic computation.

The Kalman TD model makes a number of new experimental predictions. First, it predicts that a host of post-training manipulations, identified as problematic for traditional associative learning (Gershman, 2015; Miller et al., 1995), should have systematic effects on dopamine responses. For example, extinguishing the blocking cue in a blocking paradigm causes recovery of responding to the blocked cue in a subsequent test (Blaisdell et al., 1999); the Kalman TD model predicts that this extinction procedure should cause a positive dopaminergic response to the blocked cue. A second prediction is that the OFC should exhibit dynamic cue competition and facilitation (depending on the paradigm). For example, in the sensory preconditioning paradigm (where facilitation prevails), neurons selective for one cue should be correlated with the neurons selective for another cue, such that presenting one cue will activate neurons selective for the other cue. By contrast, in a backward blocking paradigm (where competition prevails), neurons selective for different cues should be anti-correlated. Finally, OFC lesions in these same paradigms should eliminate the sensitivity of dopamine neurons to post-training manipulations.

The theory presented here does not pretend to be a complete account of dopamine; there remain numerous anomalies that will keep RL theorists busy for a long time (Dayan and Niv, 2008). The contribution of this work is to chart a new avenue for thinking about the function of dopamine in probabilistic terms, with the aim of building a bridge between RL and Bayesian approaches to learning in the brain.

## Acknowledgments

This research was supported by the NSF Collaborative Research in Computational Neuroscience (CRCNS) Program Grant IIS-120 7833.

## Footnotes

1 Preuschoff and Bossaerts (2007) made an essentially identical suggestion, but did not provide a mechanistic proposal for how the scaling term would be computed.

2 Eshel et al. (2016) have reported that dopamine neurons in the ventral tegmental area exhibit homogeneous prediction error responses that differ only in scaling. One possibility is that these neurons have different noise levels (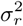) or volatility estimates (*q_d_*), which would influence the normalization term *λ_t_.*

